# Cellular signalling protrusions enable dynamic distant contacts in spinal cord neurogenesis

**DOI:** 10.1101/2024.10.11.617849

**Authors:** Joshua Hawley, Robert Lea, Veronica Biga, Nancy Papalopulu, Cerys Manning

**Affiliations:** Faculty of Biology Medicine and Health, The University of Manchester, Manchester, UK

**Keywords:** Protrusions, cytonemes, HES5, lateral inhibition, spatial pattern, Notch, spinal cord

## Abstract

In the developing mouse ventral spinal cord, HES5, a transcription factor downstream of Notch signalling, is expressed as evenly spaced clusters of high HES5-expressing neural progenitor cells along the dorsoventral axis. While Notch signalling requires direct membrane contact for its activation, we have previously shown mathematically that contact needs to extend beyond neighbouring cells for the HES5 pattern to emerge. However, the presence of cellular structures that could enable such long-distance signalling was unclear. Here, we report that cellular protrusions are present all along the apicobasal axis of individual neural progenitor cells. Through live imaging, we show that these protrusions dynamically extend and retract reaching lengths of up to ∼20μm, enough to extend membrane contact beyond adjacent cells. The Notch ligand DLL1 was found to colocalise with protrusions, further supporting the idea that Notch signalling can be transduced at a distance. The effect of protrusions on the HES5 pattern was tested by reducing the density of protrusions using the CDC42 inhibitor ML141, leading to a tendency to decrease the distance between high HES5 cell clusters. However, this tendency was not significant and leaves an open question about their role in the fine-grained organisation of neurogenesis.

## Introduction

The central nervous system’s (CNS) function depends upon spatial patterning to generate the correct cell types in the right place during embryogenesis. Previous studies have shown that morphogen gradients and transcription factor (TF) networks underlie spatial patterning by mediating the subdivision of the embryonic spinal cord in distinct but broad progenitor subdomains (Ericson et al., 1995; Briscoe et al., 2000; Cohen et al., 2013). Such subdomains generate distinct neuronal subtypes along the dorsoventral axis, for example, motor neurons ventrally and various types of interneurons more dorsally (Sagner and Briscoe, 2019).

Using a live protein reporter for the Notch target TF HES5, we have previously shown that a finer spatiotemporal organisation exists within the ventral interneuron neural progenitor domain. We previously reported that the oscillatory expression of HES5 over time is synchronised in microclusters of 3-5 cells. These microclusters are repeated along the dorsoventral axis every 2-3 cells, forming a spatially periodic pattern. Furthermore, the pattern is highly dynamic with switching of HES5 microclusters from high to low HES5 expression states on the timescale of 7 hours (Biga et al., 2021). Based on experimentation and modelling, we hypothesised that this fine-grained spatiotemporal pattern controls the rate and spacing of neurogenesis and that Notch-mediated self-organisation is sufficient to generate it (Biga et al., 2021; Hawley et al., 2022).

However, while Notch signalling is classically thought of as occurring between neighbouring cells, our modelling suggested that longer-range Notch signalling, extending further than immediate neighbouring cells, is necessary for the repetition of HES5 microclusters along the dorsoventral axis with a spatial periodicity of 3-4 cells, as summarised in Figure 1 (Biga et al., 2021; Hawley et al., 2022). Mathematically, without this long-range signalling, i.e Notch signals between immediate/adjacent neighbours, a lateral inhibition or “salt and pepper” pattern forms (2-cell spatial period shown in Figure 1). If signalling distance is extended beyond immediate neighbours this gives rise to the possibility of longer spatial periods between microclusters of cells with correlated HES5 levels and dynamics to form (Hawley et al., 2022).

**Figure 1:**
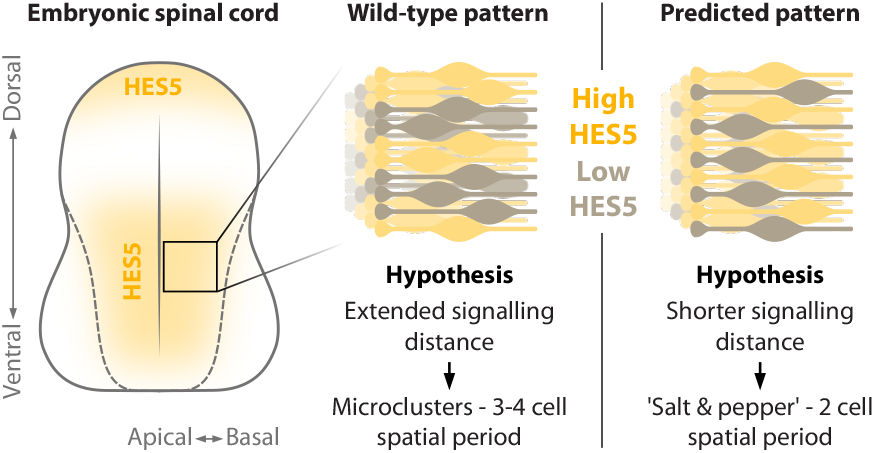
Overview of HES5 patterning in the embryonic mouse spinal cord. **Left** shows the expression regions of HES5 in a cross-sectional view of the spinal cord. **Middle** shows a magnified illustration of the wild-type HES5 expression pattern in the neuroepithelia that makes up the developing spinal cord. Yellow cells represent higher expressing cells, and grey corresponds to lower expression. In the wild-type neuroepithelia, the expression of HES5 is clustered and has a spatial period of 3-4 cells in the dorsoventral direction. This paper hypothesises that this clustered expression arises from extended Notch signalling distance between cells. **Right** shows the predicted pattern change to a ‘salt and pepper’ pattern with a shorter spatial period.

Indeed, recent findings have shown that long-range signalling events can occur via specialised cellular protrusions, such as cytonemes, that make direct contact with and signal to cells at a distance. Cytonemes are a specialised form of cellular protrusion that can exchange or present signalling molecules between cells and there is increasing evidence for cytoneme regulation of spatial patterning during embryonic development, particularly for the Wnt, Shh and Notch pathways (De Joussineau et al., 2003; Cohen et al., 2010; Rojas-Ríos et al., 2012; Bischoff et al., 2013; Nelson et al., 2013; Stanganello et al., 2015; Sagar et al., 2015; Hadjivasiliou et al., 2019; Hall et al., 2024).

In the Notch pathway, several examples of cytonemes containing Notch/Delta signalling molecules have been reported. In the Drosophila notum the regular spacing of sensory organ precursors, which give rise to mechanosensory bristles, require the presence of Notch signalling protrusions that extend beyond neighbouring cells to achieve the observed spacing between cells, and the expression of Delta promotes the formation of these protrusions (De Joussineau et al., 2003; Cohen et al., 2010; Corson et al., 2017). In developing neural tissues DLL1 containing protrusions have also been observed. In the zebrafish spinal cord, differentiating neurons transiently extend long basal protrusions in the anterior-posterior direction to deliver long-range lateral inhibition, and control the spacing of subsequent differentiation events (Hadjivasiliou et al., 2019). In mouse cortex neuroepithelia, protrusions containing DLL1 were found in both RGCs and intermediate neural progenitors, however, no functional role for these protrusions was tested (Nelson et al., 2013). In chick spinal cord, actin-dependent protrusions were observed at the apical endfeet of radial glial cells (RGCs), though their function was unclear (Kasioulis et al., 2022). Protrusions in the mouse spinal cord were recently observed to emanate from floorplate and roofplate cells carrying Shh and Wnt signals and were required for correct dorsal-ventral patterning (Hall et al., 2024).

Although cellular protrusions have been observed in the mouse spinal cord it is unclear whether they are restricted to floorplate and roofplate signalling centres and the apical endfeet of cells. The latter question is pertinent to the organisation of the neuroepithelium, where neural progenitor cells are bi-polar and elongated, extending the full apicobasal dimension of the spinal cord.

Here, we have used a live membrane reporter and sparse labelling to examine the presence of cellular protrusions in the embryonic mouse spinal cord. We report that actin-based cellular protrusions exist throughout the apicobasal dimension of the radially elongated progenitor cells. Both fixed and live spinal cord slices were imaged to assess protrusion length and we observed protrusions capable of extending contact distance beyond their immediate neighbours. We also found that these protrusions are highly dynamic with an average lifetime of 4 minutes and contain the Notch signalling ligand DELTA-LIKE 1 (DLL1). The protrusions are initiated through CDC42-dependent actin polymerisation as CDC42 inhibition by ML141 resulted in a change in protrusion density but not a reduction in protrusion length. Finally, altering the protrusion density resulted in a tendency towards smaller HES5 spatial periods (reduced by 20% as measured by Fourier transform), as predicted by our previous mathematical modelling, although the change was not statistically significant. This suggests that reducing protrusion density is not sufficient to reveal a contribution to patterning.

## Results

### Protrusions are present in neural progenitor cells of the mouse spinal cord

To examine the existence of protrusions in mouse spinal cord RGCs, we used Sox2-tamoxifen-dependent Cre-mediated (Sox2CreERT2) recombination of the membrane-EGFP membrane-tdTomato (mTmG) reporter (Figure 2A) (Snyder et al., 2013). We used a low dose of tamoxifen to achieve mosaic membrane-EGFP labelling of Sox2 positive cells, allowing individual cell membranes to be distinguished in the densely packed neuroepithelium (Figure 2B & Movie 1). The bipolar morphology of neural progenitor cells is clearly seen with cellular processes contacting both apical and basal surfaces (Figure 2B). The higher magnification images in Figure 2C-D and Movie 2 reveal the presence of thin filopodia-like protrusions on most of these elongated cells, emanating from anywhere along the apicobasal length of the cell. Protrusions were traced and are highlighted with white lines in most panels in Figure 2C-D. Therefore the mTmG Sox2CreERT2 system proved sufficient to proceed with both fixed and live imaging to measure and characterise protrusions in mouse neuroepithelia.

**Figure 2:**
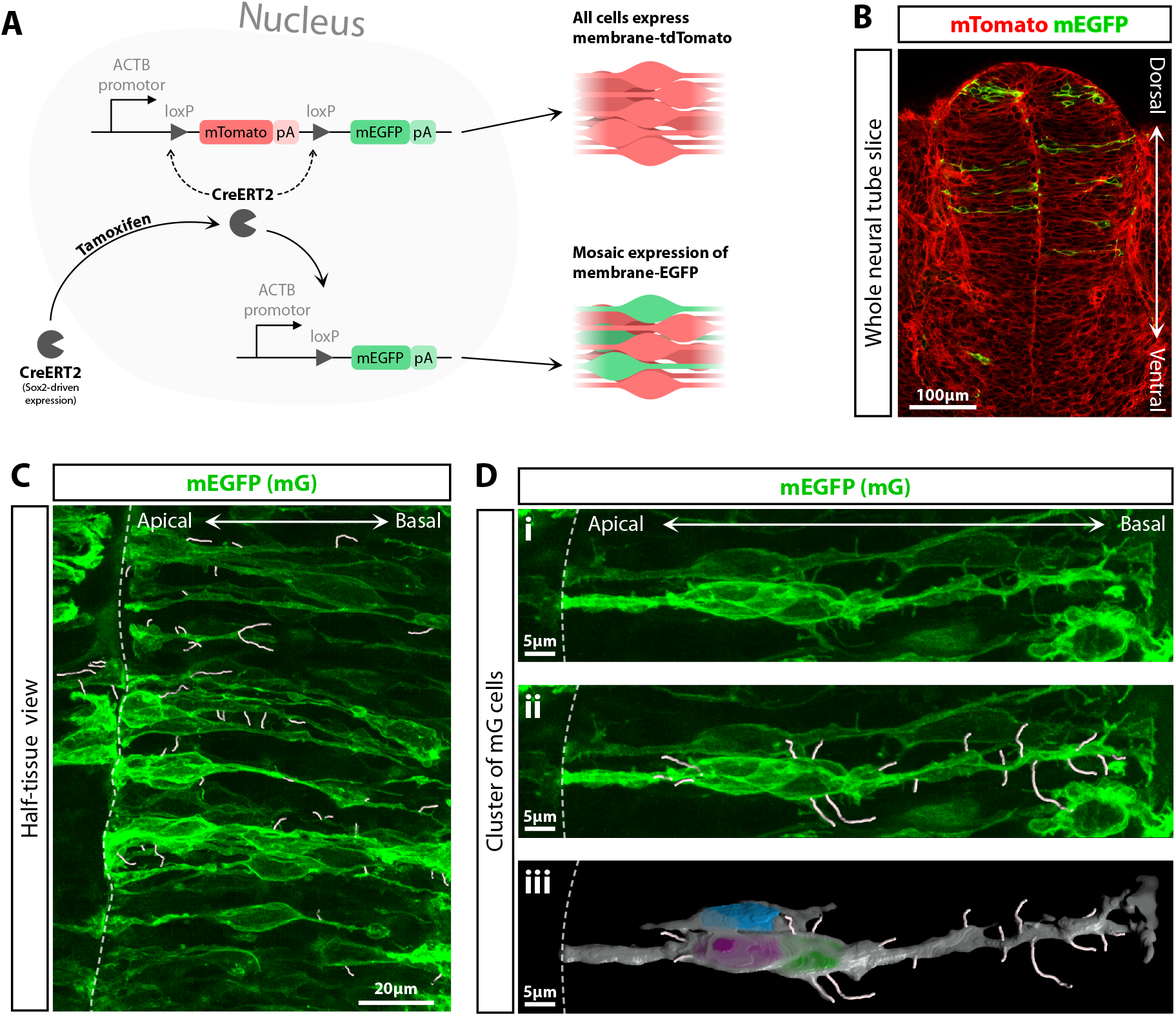
The Sox2CreERT2 mTmG system is used to mosaically label the membranes of neural progenitor cells in the spinal cord. **A** Schematic of the Sox2CreERT2 mTmG system: CreERT2 refers to Cre recombinase which targets and excises genes located between two loxP sites and also has an estrogen ligand-binding domain that requires tamoxifen for it to be localised to the nucleus. The ACTB promotor ubiquitously expresses the downstream genes. pA is the abbreviation for polyA tails. mTomato is membrane-targeted tdTomato fluorophore, and mEGFP is membrane-targeted EGFP. See Materials & Methods 1.1 for further details. **B-D** Confocal images of fixed Sox2CreERT2^+/−^ mTmG^+/−^ E10.5 mouse embryos (Materials & Methods 1.1 & 1.2). **B** Single z-slice image of a whole spinal cord slice showing membrane-tdTomato (mT) in red and tamoxifen-induced membrane-EGFP (mG) in green (see also Movie 1). **C** A z-projection of one side of a spinal cord slice with protrusions traced in white (Materials & Methods 1.5). **D** Shows a magnified view of a group of RGCs (see also Movie 2). **D i** Shows the raw data, **D ii** shows the data with traced protrusions in white. **D iii** shows a 3D fill of the membrane (grey) and nuclei of the cells (blue, magenta, green). Separate embryos were used to generate each panel image.

### Protrusion lengths and angles in fixed tissue

To understand how far protrusions extend the contact distance in RGCs, both fixed and live spinal cord slices were used to characterise protrusion lengths, angles, and temporal dynamics (see Materials & Methods 1.5). In fixed tissue, a range of protrusion lengths was observed (Figure 3A) with the average protrusion length being 3.4 ± 3.3μm (median ± IQR). Figure 3A shows a skewed length distribution with some protrusions exceeding 20μm.

**Figure 3:**
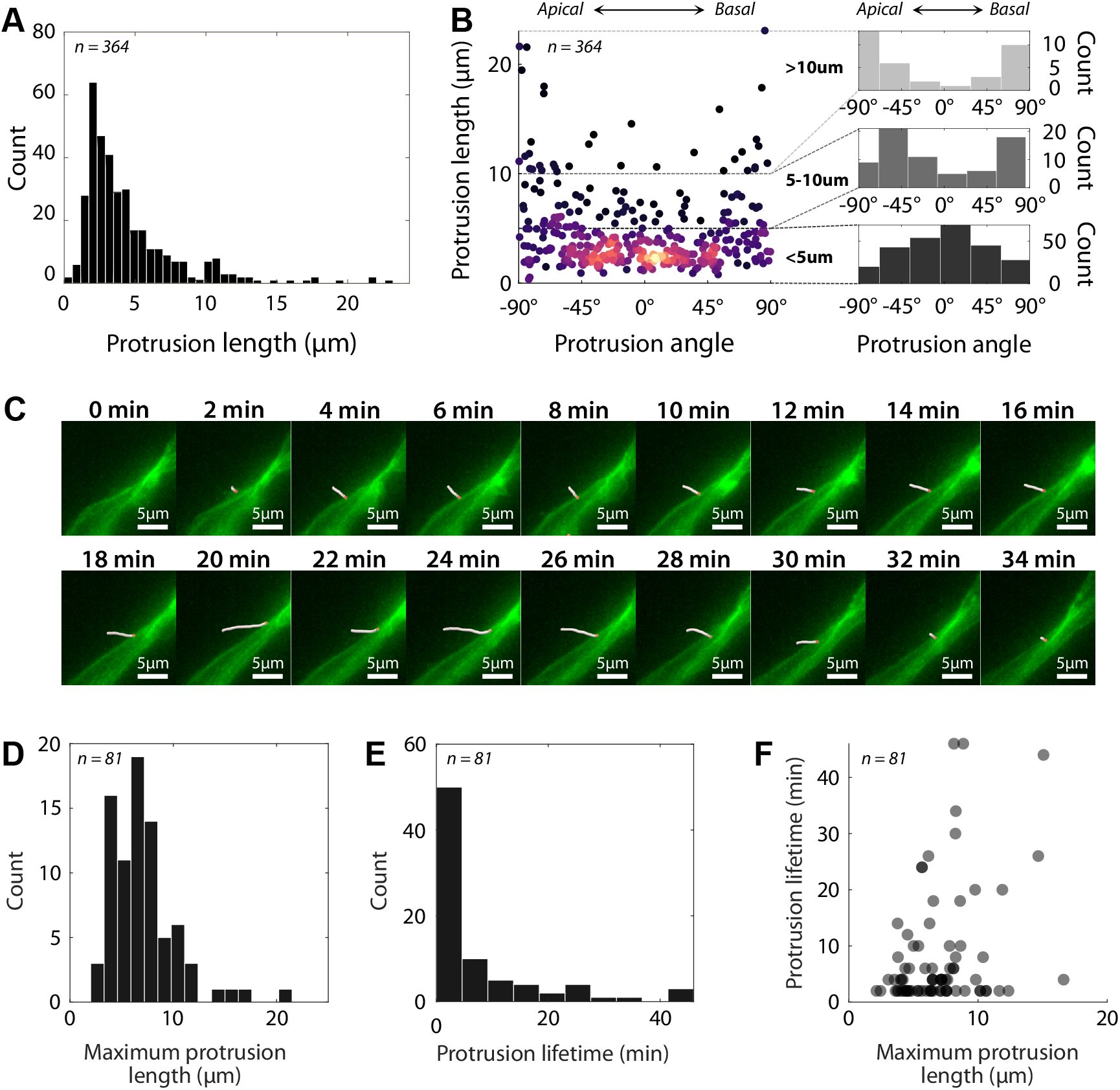
Quantitative characterisation of protrusion length, angle, and dynamics in both fixed and live tissue. **A** Histogram of protrusion lengths measured in 12 fixed slices from 5 different E10.5 Sox2CreERT2^+/−^ mTmG^+/−^ embryos (n=364 protrusions). **B** Angle of the protrusions relative to the apicobasal axis (using the same data set as in **A**). The colouring of the points indicates plot density with lighter colours being higher density. 0° indicates a protrusion oriented perpendicular to the apicobasal axis, −90° is a protrusion oriented towards the apical surface and 90° is a basally directed protrusion. The smaller histogram plots to the right show how the distribution varies with protrusion length. **C** Frames from live imaging (image captured every 2 minutes) showing the extension and retraction of a protrusion traced in white (the red dot indicates where the protrusion joins the cell body). Movie 3 shows a timelapse with tracked protrusions. **D-F** Characterisation of live-imaged protrusions (n=81 protrusions). A protrusion with an outlier lifetime of 150min was excluded from the data shown. **D** Distribution of maximum protrusion lengths measured for each protrusion tracked in time-lapse imaging. Average maximum length was 6.5 ± 3.6μm (median ± IQR). **E** Distribution of protrusion lifetimes (excluding one outlier at 150min lifetime). Average lifetime was 4 ± 7 min (median ± IQR). **F** Maximum protrusion length plotted against protrusion lifetime.

Next, protrusion angles were measured relative to the apicobasal axis where −90° is defined as pointing apically, +90° is pointing basally, and 0° pointing perpendicular from the apicobasal axis (along the dorsoventral axis). Shorter-length protrusions were more likely to be perpendicular to the apicobasal axis, which can be seen in Figure 3B as a higher density of angles around 0° for protrusions less than 5μm. In the 5-10μm and *>*10μm range, the distribution becomes spread out, becoming more aligned with the apicobasal axis with increasing length.

The persistence length of filopodia, which is the length at which a polymer such as actin will start to bend, is on the order of 10μm (Mogilner and Rubinstein, 2005; Wisanpitayakorn et al., 2022). This persistence length, combined with the fact the neuroepithelia is a densely packed tissue, suggests that filopodia may be mostly forming perpendicular to the apicobasal axis, and then bending as they lengthen and are obstructed by neighbouring cells. Indeed, many curved and kinked longer protrusions can be seen in Figure 2C&D.

### Protrusion dynamics in live tissue

To characterise the temporal dynamics of protrusions we used live imaging of E10.5 *ex vivo* mouse spinal cord slices embedded in collagen (Materials & Methods 1.4). We traced individual protrusions at every time step (every 2min), as shown in Figure 3C and Movie 3. From this, we obtained the maximum extension length and average lifetime (Figure 3D&E). The average maximum extension length was found to be 6.5 ± 3.6μm, and the average lifetime was 4 ± 7min (median ± IQR). Compared to measurements in fixed tissue, the maximum protrusion length distribution is shifted towards longer lengths, and there is a tendency for longer protrusions to have a longer lifetime (Figure 3F). Overall, live imaging allows for a more accurate estimate of how far protrusions can reach and reveals that these protrusions are highly dynamic in their extension and retraction.

### Protrusions can extend contact distance beyond immediate neighbours

We next wanted to put the measured length distributions into the context of neighbouring and non-neighbouring cells. To understand what sort of distance a protrusion would need to traverse to reach a non-neighbouring cell, we characterised the morphology of RGCs (Figure 4A). This was done by measuring three distinct parts of the cell: the apical endfoot width (5.2 ± 1.8μm, mean ± SD), the process width (1.6 ± 0.9μm), and the cell body width (7.7 ± 1.1μm).

**Figure 4:**
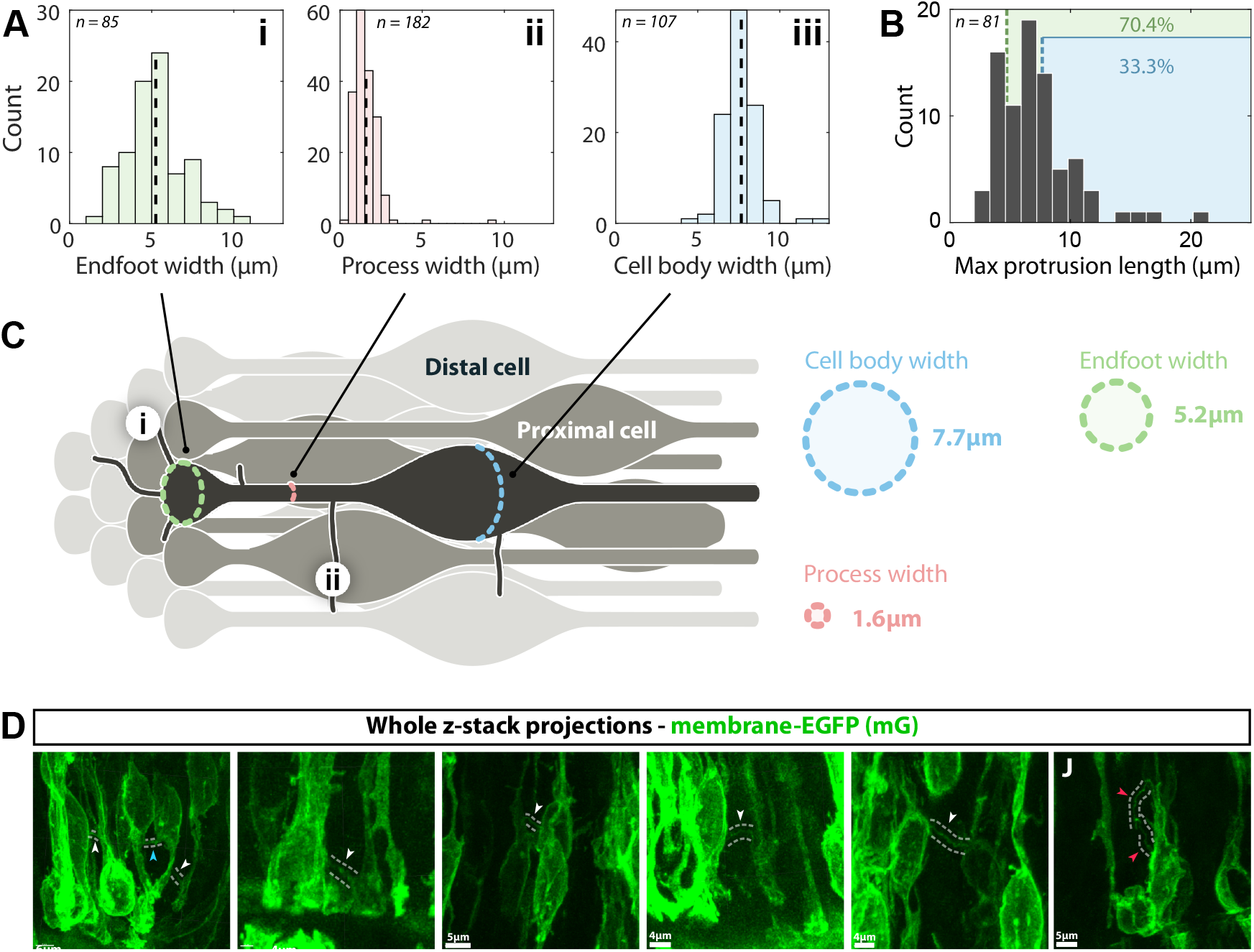
**A** Width measurements of different parts of RGCs. **A i** Histogram of apical endfoot widths (n=85, average width 5.2 ± 1.8μm (mean ± SD)). **A ii** Histogram of RG process widths (n=182, average width 1.6 ± 0.9μm (mean±SD)). **A iii** Histogram of cell body widths (n=107, average width 7.7 ± 1.1μm (mean ± SD)). See Materials & Methods 1.6 for how protrusions and cell widths were measured. **B** The maximum protrusion length histogram from Figure 3D (n=81), but with the mean measured RGC widths highlighted in different colours. Green is the mean endfoot width (70.4% of protrusions are longer than this), and blue is the mean cell body width (33.3% of protrusions are longer than this). **C** Idealised arrangement of RGCs scaled by the measured cell widths. The cell of interest is labelled in dark grey, proximal cells in mid-grey, and distal cells in light grey. **C i** A protrusion with a length equal to the mean apical endfoot width, emanating from an apical endfoot. **C ii** highlights a protrusion with a length equal to the mean RGC cell body width, emanating from a process. **D** Examples of protrusions enabling contacts between cells marked with membrane-EGFP. Dashed lines highlight protrusions. White arrowheads indicate distal cells making contact via a protrusion. The blue arrowhead highlights proximal cells making contact via a protrusion. Red arrowheads show an example of two protrusions originating from the same cell contacting each other.

From these measurements, we constructed a to-scale ‘idealised’ schematic of how RGCs are packed by using the mean width measurements (Figure 4C). A main assumption in the schematic is that the packing of the apical endfeet determines how close together the cells are. Additionally, RGCs are not perfectly straight in the tissue; they can curve and potentially contact new neighbours along their apicobasal length. This likely results in changes in their neighbourhood along their apicobasal axis, as shown in mouse lung epithelia (Gomez et al., 2021). However, to simplify the interpretation of the protrusion lengths and what is defined as a neighbouring/non-neighbouring cell, we assume that RGCs are straight and don’t intercalate. In Figure 4C, we refer to neighbouring cells as *proximal* cells and they are defined as cells whose apical endfeet directly contact the cell of interest (dark grey RGC). Non-neighbouring cells are referred to as *distal* cells.

The location of protrusions on an RGC determines whether a distal cell will be contacted. For example in Figure 4Ci, if a protrusion emanates from the apical endfoot, it would need to be longer than 5.2μm. In this case, 70.4% of protrusions would be long enough to cover this distance (using the maximum extension lengths shown and highlighted in green in Figure 4B). Alternatively in Figure 4Cii, if a protrusion emanates from a radial process and needs to cross the width of a cell body (7.7μm), then 33.3% of protrusions would contact distal cells.

Therefore protrusions are expected to be capable of making distal contacts, and these were indeed observed in Sox2CreERT2 mTmG embryos (Figure 4D). The idealised model highlights how the likelihood of making a distal contact depends on where protrusions originate on an RGC.

### DLL1 is present in neural progenitor protrusions

Next, we carried out immunofluorescent (IF) staining in mTmG Sox2CreERT2 spinal cord slices to see if protrusions contain the Notch ligand DLL1, as this would be required for the distal contacts to mediate Notch signalling.

Figure 5Ai shows an IF stain for DLL1 (Materials & Methods 1.2). As previously reported, two bands where no DLL1 is expressed, corresponding to the p1, d6 progenitor domains, were observed (indicated by the white arrowheads) (Marklund et al., 2010). At higher magnification, punctate expression was seen, largely localised to the plasma membrane (Figure 5Aii). When individual neural progenitor cells were visualised in Sox2CreERT2 mTmG embryos, DLL1 localisation within protrusions was observed (Figure 5Bi-iv), indicating the possibility that Notch signalling is being transduced via these cellular structures.

**Figure 5:**
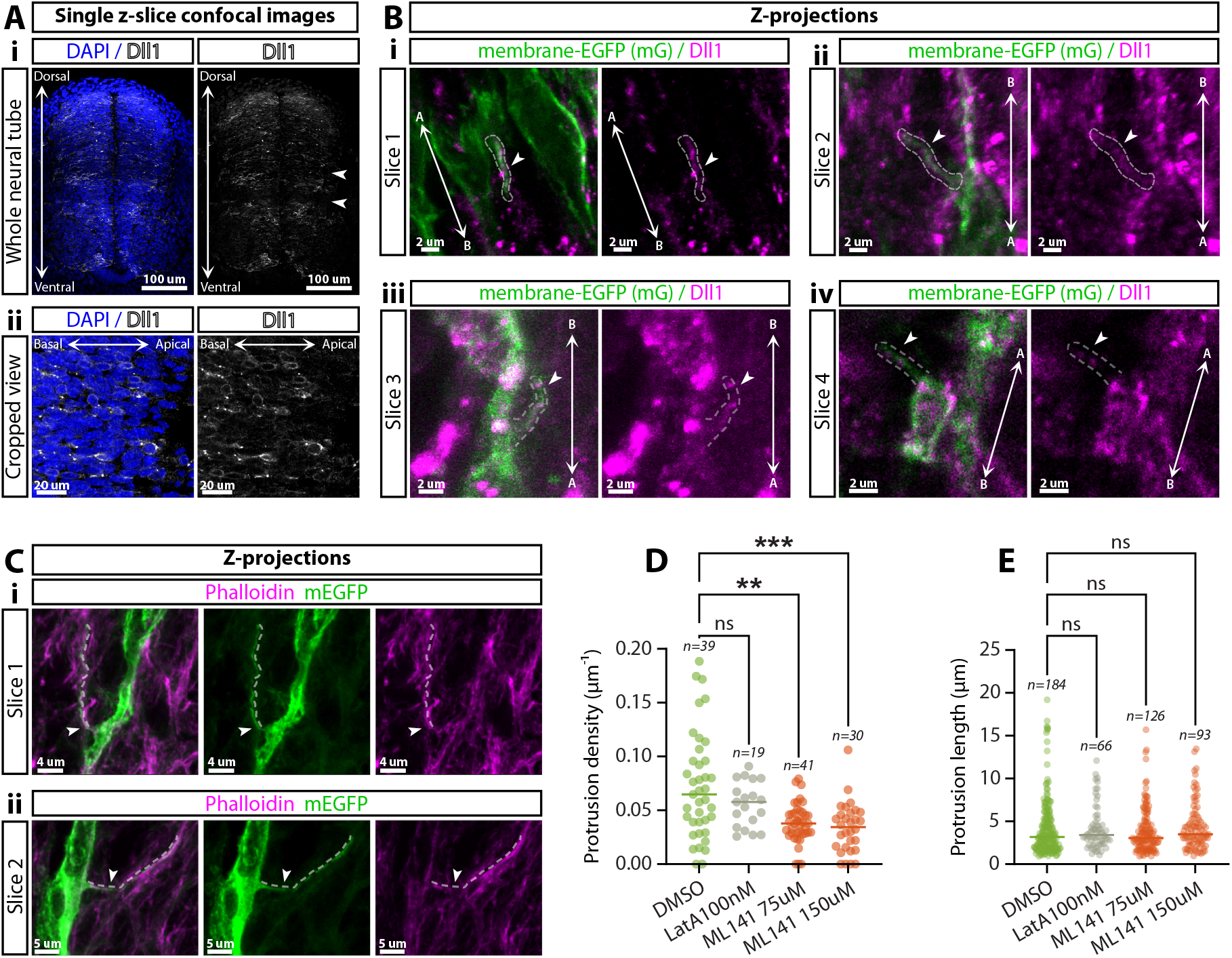
IF staining in E10.5 mTmG^+/−^ Sox2CreERT2^+/−^ cryosections (Materials & Methods 1.2). **A i** Single z-slice confocal image of a whole spinal cord cryosection showing DAPI and DLL1 immunostaining. The two white arrowheads highlight low expression bands of DLL1 where other ligands are expressed as reported in (Marklund et al., 2010). **A ii** Higher magnification images of DAPI and DLL1. **B i-iv** Z-projections of mEGFP and DLL1 where the dashed lines and white arrowheads highlight protrusions that contain DLL1. Double arrows marked with A and B indicate the apicobasal axis. **C i&ii** Co-localisation of phalloidin in mEGFP marked protrusions (arrows and dashed lines highlight the protrusion). **D&E** Embryos were sliced on a vibratome as in Materials & Methods 1.3, and left for 1h in DMSO, Latruculin A, or ML141, and then fixed in PFA for imaging. **D** Protrusion density measured in fixed tissue after 1h of treatment in either DMSO (n=39), Latrunculin A 100nM (n=19), ML141 75uM (n=41), or ML141 150uM (n=30) (n-number indicates the number of cells density was measured for) (Materials & Methods 1.5). The statistical test used was a two-tailed Mann-Whitney U test. P-values from left to right: >0.99, 0.0019, 0.0003. **E** Protrusion length measured in fixed tissue after 1h of treatment in either DMSO (n=184), Latrunculin A 100nM (n=66), ML141 75uM (n=126), or ML141 150uM (n=93) (n-number indicates the number of protrusions measured) (Materials & Methods 1.5). The statistical test used was a two-tailed Mann-Whitney U test. P-values from left to right: >0.99, >0.99, 0.53.

### Protrusions are actin-based and are reduced in density by the CDC42 inhibitor ML141

Signalling cytonemes, a specialised form of filopodia-like protrusions, can be generated through either a cell dislodgement mechanism, where thin tubes form after adjacent cells touch and move apart; or an actin driven protrusion mechanism. Our live imaging showed active extension and retraction of protrusions indicating an actin-driven protrusion mechanism. To confirm that the protrusions were actin-based we used phalloidin staining. As expected, we observed F-actin enrichment at the apical surface of the mouse spinal cord, but F-actin was also present in protrusions (Figure 5Ci&ii).

Previous work has shown that actin-based cytonemes are initiated by cytoskeletal regulators such as CDC42 (Watson et al., 2017). Therefore we tried to alter the length of these protrusions by adding Latrunculin A, to inhibit actin polymerisation and ML141 to inhibit CDC42 in *ex vivo* embryonic spinal cord slices. Low concentrations of Latrunculin A that did not lead to whole tissue disruption had no effect on the protrusion length or protrusion density, as shown in Figure 5D&E. However inhibition of CDC42 using ML141 led to a significant reduction in protrusion density (Figure 5D) as previously reported in other tissues (Fantin et al., 2015; Grönloh et al., 2023), but protrusion length was unchanged (Figure 5E). Taken together, this suggests that RhoGTPase CDC42-dependent actin dynamics initiates protrusion formation in mouse spinal cord neural progenitor cells but does not contribute to protrusion extension.

### Reduction in protrusion density does not significantly alter the HES5 spatial period

Our previous modelling suggested that an extended Notch signalling distance, potentially through protrusions, generated spatially periodic HES5 microclusters along the dorsoventral axis (Hawley et al., 2022). The model predicted that reducing the signalling distance would lead to a more lateral inhibition-like high-low-high HES5 pattern in adjacent cells and hence a shorter HES5 spatial period. Therefore we experimentally tested whether the decrease in protrusion density caused by the addition of the CDC42 inhibitor ML141 would lead to a reduction in the spatial period of HES5 expression in the spinal cord.

spinal cord slices from embryos with endogenously tagged Venus::HES5 were incubated either in DMSO (control) or 75uM ML141 for 8h on an air-media interface and then imaged using a confocal microscope (Materials & Methods 1.3). Slice quality was controlled for, and the left and right sides of the spinal cord were used to extract fluorescence intensity along the dorsoventral axis. This was followed by Gaussian blurring to remove the influence of periodicity from the gaps between nuclei (10μm periodicity). Detrending was applied to the mean signal to remove the effect of the HES5 domain edges. Finally, the spatial period was estimated either by use of a Fast Fourier Transform (FFT) or autocorrelation function (ACF). An overview of the spatial period analysis pipeline is shown in Figure 6A&B – full details are given in Materials & Methods 1.7.

**Figure 6:**
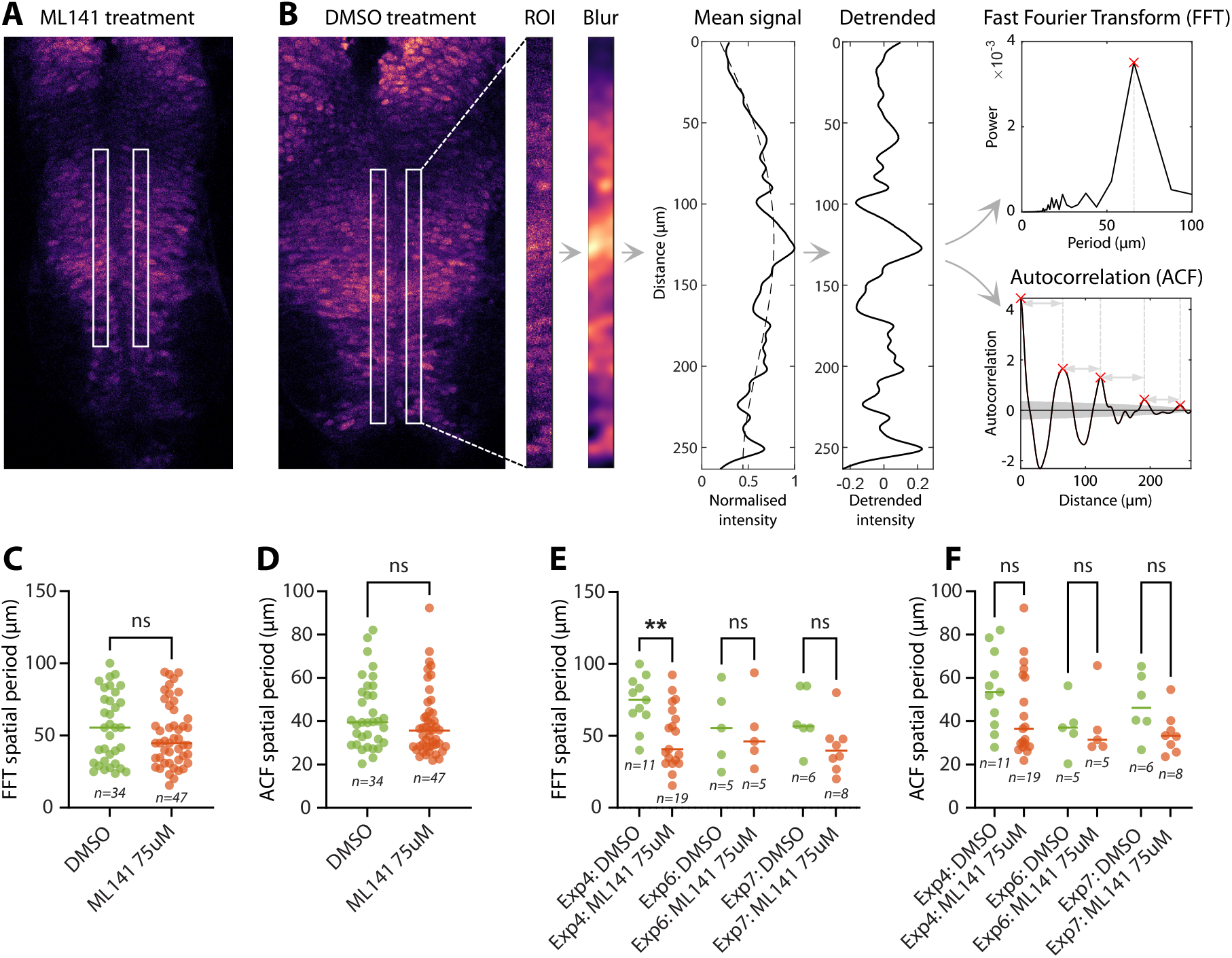
Spatial periodicity in HES5 expression pattern after DMSO or ML141 treatment. **A** Venus::HES5 fluorescence in an ML141-treated spinal cord slice (Materials & Methods 1.1). White boxes indicate the regions of interest (ROI) used for spatial period analysis. **B** Analysis pipeline for determining HES5 spatial periodicity along the dorsoventral axis of the spinal cord within the HES5 p0-p2 and pMN domains. The analysis starts with a single z-slice image taken from a confocal z-stack. Next, ROIs are drawn on either side of the ventricle. Then a Gaussian blur with a standard deviation of 3μm is applied to remove the effects of internuclear periodicity (10μm). The mean signal is extracted for each row of the ROI, and a polynomial of order 4 (dashed line) is used to detrend. An FFT (Fast Fourier Transform) and ACF (autocorrelation function) method is then used to process the detrended signal. The red cross in the FFT indicates the dominant period and the red crosses in the ACF indicate peaks above the bootstrapping significance threshold visualised as the grey area. Distances between peaks (grey arrows) in ACF are averaged to produce the final spatial period. See Materials & Methods 1.7 for full details of the pipeline. **C-F** Spatial periods of Venus::HES5 fluorescence, where each dot corresponds to an individual ROI from a spinal cord slice. A two-tailed Mann-Whitney U test was used to determine statistical differences in the distributions in all graphs. **C** FFT spatial period of all experimental repeats combined (median values are Med_DMSO_ = 55.4μm (n=34), Med_ML141_ = 44.7μm (n=47), p_val_ = 0.54). **D** ACF spatial period of combined experimental repeats (median values are Med_DMSO_ = 39.6μm (n=34), Med_ML141_ = 35.7μm (n=47), p_val_= 0.14). **E** FFT spatial period of individual experiments that have ≥5 data points - enough for the statistical test (Med_Exp4:DMSO_ = 75.1μm (n=11), Med_Exp4:ML141_ = 40.7μm (n=19), p_val_= 0.0053; Med_Exp6:DMSO_ = 55.4μm (n=5), Med_Exp6:ML141_ = 46.2μm (n=5), p_val_ > 0.99; Med_Exp7:DMSO_ = 56.6μm (n=6), Med_Exp7:ML141_ = 39.6μm (n=8), p_val_= 0.0057). **F** ACF spatial period of individual experiments that have ≥5 data points - enough for the statistical test (Med_Exp4:DMSO_ = 53.4μm (n=11), Med_Exp4:ML141_ = 36.6μm (n=19), p_val_ = 0.10; Med_Exp6:DMSO_ = 37.0μm (n=5), Med_Exp6:ML141_ = 31.5μm (n=5), p_val_=0.69; Med_Exp7:DMSO_ = 46.2μm (n=6), Med_Exp7:ML141_ = 33.2μm (n=8), p_val_ = 0.081).

Figure 6C&D show the combined spatial periods from all experimental repeats as calculated by the FFT and ACF methods respectively. The FFT spatial period in Figure 6C went from a median of 55.4μm in DMSO to 44.7μm in ML141. Likewise, the ACF spatial period in Figure 6D went from a median of 39.6μm in DMSO to 35.7μm in ML141. Neither of these decreases were found to be significantly different. We also examined individual experiment spatial period changes in Figure 6E&F. This again showed the same tendency for the ML141 condition to have a smaller spatial period compared to DMSO, however only one experiment showed a significant decrease. In summary the experimentally measured spatial period of HES5 along the dorsoventral axis had a tendency to decrease when protrusion density was decreased, which agreed with our previous modelling predictions.

## Discussion

Thin cellular extensions are of increasing importance in cell signalling; they have been described to contribute to many of the main developmental signalling pathways such as Notch (De Joussineau et al., 2003; Cohen et al., 2010; Nelson et al., 2013; Hadjivasiliou et al., 2019), Wnt (Stanganello et al., 2015; Sagar et al., 2015; Hall et al., 2024; Zhang et al., 2024), Hedgehog (Rojas-Ríos et al., 2012; Bischoff et al., 2013; Hall et al., 2024), BMP (Hsiung et al., 2005; Inaba et al., 2015; Wilcockson and Ashe, 2019), and FGF (Du et al., 2018).

Here, we have interrogated the presence of cellular protrusions along the apicobasal axis of mouse spinal cord RGCs with a view of testing their potential role in extending Notch signalling distance and driving Notch-dependent gene expression patterning. We have found that such protrusions can reach up to ∼20μm with a median length of 6.5 ± 3.6μm (median ± IQR). We found that protrusions can extend contacts beyond neighbouring cells, but that the likelihood of this depends upon where protrusions emanate from on a given RGC. Protrusions are also highly dynamic in their extension and retraction with an average lifetime of 4 ± 7 min (median ± IQR). While previous findings showed the presence of cellular protrusions in the apical endfeet (Kasioulis et al., 2022) our findings show that they are a lot more extensive as they cover the extent of RGCs along the apicobasal axis.

The protrusions we observed have shared cellular features with cytonemes, a specialised form of filopodia, as they are thin actin-based structures as shown by phalloidin staining and their sensitivity to CDC42 inhibition and contain the signalling ligand DLL1. We describe a CDC42-dependent mechanism for protrusion generation, in contrast to the developing chick limb bud where CDC42 perturbations were not effective in perturbing mesenchymal filopodia formation ((Sanders et al., 2013)). This indicates that mechanisms driving protrusion and cytoneme formation may be tissue-specific. MyoX is downstream of CDC42 (Bohil et al., 2006) and has recently been shown to regulate initiation of protrusions in mouse spinal cord (Hall et al., 2024). Interestingly MyoX mutants have disrupted Notch signalling as identified by RNAseq (Hall et al., 2024). Further work is required to determine which aspects of the pathway are dysregulated and whether this includes Notch-dependent spatial patterning.

While cytonemes can be involved in local cell-cell signalling, longer-range protrusions which contact non-neighbouring cells can give rise to more complex patterns akin to Turing morphogen reaction-diffusion patterns (Hadjivasiliou et al., 2016; Vasilopoulos and Painter, 2016; Bajpai et al., 2021; Hawley et al., 2022). We hypothesised that protrusions mediate non-neighbouring contacts and we subsequently imaged examples of protrusions extending contact distance beyond immediate neighbours. Further, we found that where protrusions emanate from on RGCs determines how likely non-neighbouring contacts are due to the varying distances they need to traverse.

With few exceptions, the function of protrusions or cytonemes is enigmatic. For example, Kasioulis et al. also inhibited protrusions using truncated WAVE1–eGFP in the chick spinal cord and found that while differentiating neurons reduce their protrusion length it did not affect the number of differentiating cells (Kasioulis et al., 2022). They also found that inhibition of these protrusions did not disrupt neuroepithelium integrity or apical adherens junctions. Our previous mathematical modelling suggested that distant cell contacts mediate the regular spacing of HES5 microclusters with related levels of expression, along the dorsoventral axis of the spinal cord. We furthermore predicted that this spatial HES5 organisation would help even out differentiation events in space by extending Notch/Delta signalling beyond neighbouring cells. Indeed, the presence of DLL1 within the cytonemes reported here suggests that Notch/Delta signalling may occur in these longer-range protrusions. In our mathematical model, the distance between microclusters in the spatially periodic pattern of HES5 is dependent on longer range Notch signalling (J R Soc Interface, 2022). Without this long range signalling, the spatial HES5 expression would revert to the classical lateral inhibition/”salt and pepper” Notch patterning (alternating high and low expressing cells) (Hawley et al., 2022). To probe this prediction and test potential functionality, we perturbed the formation of protrusions and looked at the resulting spatial pattern of HES5.

We initially aimed to alter protrusion length through the actin-disrupting agent Latrunculin A, but this blunt approach led to changes in tissue morphology making it difficult to address spatial patterns of gene expression. At reduced concentrations that maintained tissue architecture, no effect on protrusion length was observed. Therefore, we turned to the CDC42-inhibitor ML141, a more specific inhibitor of actin-based protrusion. This perturbed protrusion density without changing protrusion length, and ultimately we found that lower protrusion density led to a tendency to decrease the spatial period of HES5. However, although the mean spatial period was consistently reduced, the reduction was not statistically significant.

One potential explanation relates to the observation that our experiments ended up altering protrusion density and not protrusion length. It is possible that density changes may not lead to the same changes in spatial patterning as the predicted changes from protrusion length perturbation. Interestingly the Notch-Delta protrusion model constructed by Cohen et al. found that spatial periodicity of Delta-expressing cells is robust to changes in protrusion signalling likelihood when protrusions are dynamic. If we interpret the signalling likelihood in that model to be analogous to protrusion density, their work may explain why no significant change in the HES5 spatial period was observed when perturbing only protrusion density. More specific perturbations to protrusion length will be required to fully determine if protrusions play a role in the spatial patterning and spatial period of HES5 expression.

An alternative mechanism that could extend cell-cell contact distance is the intercalation of RGCs, which we did not include in our idealised neuroepithelial model. In pseudostratified mouse lung epithelia, Gomez et al. found that cells change neighbours along the apicobasal length due to curvature and changes in cell width (Gomez et al., 2021). This resulted in a broader neighbourhood of cells than expected by looking at the apical surface contacts. Further work in the spinal cord could define the extent of a cell neighbourhood using a multicolour labelling system such as Brainbow (Dumas et al., 2022; Malaguti et al., 2024) or more agnostic approaches to understand the extent of cell-cell contacts such as the synNotch system (Zhang et al., 2022; Malaguti et al., 2022).

## Acknowledgements

The authors thank Raman Das and his lab for help with live imaging and access to microscopes and the University of Manchester Bioimaging core facility.

## Data & Code Availability

All code for processing data and the image analysis pipeline is written in MATLAB. Statistical analysis was carried out in GraphPad Prism. All code and data is available on GitHub: https://github.com/Papalopulu-Lab/Hawley2024.

## Materials & Methods

### 1.1 Mouse lines and Tamoxifen administration

Animal experiments were performed within the conditions of the Animal (Scientific Procedures) Act 1986. mTmG (B6.129(Cg)-Gt(ROSA)26Sor^tm4(ACTB-tdTomato,-eGFP)Luo^/J homozygous, strain number 007676) and Sox2CreERT2 (B6;129S-Sox2^tm1(cre/ERT2)Hoch^/J heterozygous strain number 017593) were obtained from The Jackson Laboratory (JAX). Venus::HES5 knock-in mice (ICR.Cg-Hes5<tm1(venus)Imayo>) (Imayoshi et al., 2013) were obtained from Riken Biological Resource Centre, Japan. In these mice the mVenus fluorescent protein is fused to the N-terminus of endogenous HES5. E0.5 was considered as midday on the day a plug was detected. Pregnant mTmG Sox2CreERT2 females were injected with 250μg or 125μg (for more sparse cellular labelling) of tamoxifen in 100ul of corn oil 15-20h before harvesting the embryos at E10.5. mTmG Sox2CreERT2 embryos of the correct genotype were screened based on the detection of mosaic EGFP expression in the head.

### 1.2 Immunofluorescence staining

Trunks of E10.5 embryos for cryo-sectioning were fixed in 4% PFA for 1 hour at 4°C, followed by 3 quick washes with 1xPBS and 1 longer wash for 1 hour at 4°C. Embryos were equilibrated overnight in 30% sucrose (MERCK) at 4°C before mounting in Tissue-Tek OCT (Sakura) in cryomoulds and freezing at −80°C. 20-40μm sections were cut on Leica CM3050S cryostat. Sections were washed twice with PBS. Sections were then blocked in 10% Dako serum-free blocking reagent (Agilent) for 15 minutes followed by incubation in primary antibody diluted in blocking reagent for 2h at room temperature or overnight at 4°C. Sections were washed 3*×*10 minute PBS + 0.05% Tween-20 (MERCK) washes. Fluorescent Alexafluor-conjugated secondary antibodies were incubated for 1 h at room temperature. Sections were washed 3*×*10 minute PBS + 0.05% Tween-20, then incubated with DAPI (ThermoFisher Scientific) and 647-Phalloidin (ThermoFisher Scientific, 1:500) in PBS 5-15 minutes. Sections were washed twice with PBS and once with H2O before mounting with Moviol 4-88 (MERCK). Primary antibody: sheep anti-DLL1, 1:100 (R&D Systems AF5026). Secondary antibody: Alexafluor 647 anti-sheep, 1:500 (ThermoFisher Scientific). Images were acquired using a Zeiss LSM 880 with a 40× 1.30 NA objective.

### 1.3 Embryo slicing and culture

E10.5 mTmG Sox2CreERT2 embryos were embedded in 4% low-gelling temperature agarose (MERCK) containing 5mg/ml glucose (MERCK). 200μm transverse slices of the trunk around the forelimb region were obtained with the Leica VT1000S vibratome and released from the agarose. Embryo and slice manipulation was performed in phenol-red free L-15 media (ThermoFisher Scientific) on ice and the vibratome slicing was performed in chilled 1xPBS (ThermoFisher Scientific). Spinal cord slices were cultured on a 12mm Millicell cell culture insert (MERCK) with DMEM F-12 media(ThermoFisher Scientific) containing 4.5mg/ml glucose, 1x MEM non-essential amino acids (ThermoFisher Scientific), 120μg/ml Bovine Album Fraction V (ThermoFisher Scientific), 55μM 2-mercaptoethanol, 1x GlutaMAX (ThermoFisher Scientific), 0.5x B27 (ThermoFisher Scientific) and 0.5x N2 (ThermoFisher Scientific) and incubated at 37°C and 5% CO2. Slices were cultured with DMSO or ML141 (MedChemExpress) and fixed in 4% PFA for 1hr. Images were acquired using a Zeiss LSM 880 with a 40× 1.30 NA objective.

### 1.4 Live imaging of spinal cord slices

Spinal cord slices were embedded in type-1 Collagen (Corning) on a glass-bottomed dish (Greiner Bio-one) as described previously (Das et al., 2012). Slices were imaged using a Zeiss Cell Observer Z1 with an environment chamber maintained at 37°C and 5% CO2. Images were acquired with a 40× 1.2 NA silicone immersion objective (Carl Zeiss), a Colibri 7 light-emitting diode (LED) light source (Carl Zeiss) and a Flash4 v2 sCMOS camera (Hamamatsu). Image stacks were acquired using minimal exposure times (20 to 50 ms each channel) using intervals of x min between exposures.

### 1.5 Measuring protrusion length, density, and dynamics

The filament tool was utilised in Imaris (Oxford Instruments) in a semiautomatic way whereby automatic filament tracking was skipped and instead, the beginning and end point of the protrusion were defined manually and then the software identified the path of brightest pixels between those two points. This method worked well for sufficiently bright protrusions, and protrusion length was measured as the summed length of each tracked filament. To measure the angle of the protrusions, a reference frame where the y-axis was lined up along the ventricle of the spinal cord, and the x-axis was aligned to the apicobasal direction. From this, the angle from the start and end point was calculated relative to the apicobasal axis and depending on whether it was left or right of the ventricle, angles were corrected so that −90° was towards the ventricle (apical-pointing) and +90° was basal-pointing.

To measure protrusion density, the length of each RGC that was distinguishable from its neighbours was traced, then the number of protrusions associated with that cell was divided by the length of the RGC to get protrusions per μm length of the cell.

### 1.6 Cell width measurements

The measurement tool in Imaris (Oxford Instruments) was used to determine the width of apical endfeet, processes, and cell bodies. For the apical endfeet, the widest part of the endfoot bulge in the apicobasal direction was measured. The width of processes varies along the apicobasal axis, and so multiple measurements were made at intervals of roughly every 10μm along the apicobasal axis, meaning each cell contributed multiple process width measurements to the data set. Measurements were only made of processes that were distinguishable from other cells marked with mG. Finally, the cell body width was measured at the widest point perpendicular to the apicobasal axis.

### 1.7 Spatial period analysis pipeline

Images were processed using Matlab (code and data available here). Each spinal cord Venus::HES5 image was composed of multiple z-stacks. From each image, an ROI was extracted at every 10μm apart in the z-depth until the intensity became too low to distinguish Venus fluorescence from background easily. Once these images were loaded into Matlab, boxes 15μm wide were drawn along the

Regions of interest (ROI) of 15μm were drawn along the dorsoventral axis as close to the ventricle on each side of the ventricle. The quality of slices was determined by visual inspection of the ROI; sufficiently bright venus signal compared to the background signal and no gaps from the presence of blood vessels or damaged tissue.

Next Gaussian blurring was applied to the ROI signal using a standard deviation of 3μm. This value was chosen as it was sufficient to remove any spatial periodicity less than 10μm which is the distance between neighbouring cells (Biga et al., 2021). The calibration curve for the Gaussian blur standard deviation is shown in Figure S1.

The mean signal was then taken per row (in the apicobasal direction) to produce an average dorsoventral signal. This was followed by detrending using polynomial detrending with a polynomial value of 4 (inbuilt polyval and polyfit functions in Matlab).

Finally, the detrended signal was run through either a FFT transform or ACF methods. The FFT used the fft function in Matlab to produce power spectra, the peak period (the frequency with the highest power) was selected as the reported spatial period for that slice. A Fisher g-test was also carried out on each peak period to ensure it was statistically valid (as previously used in (Hawley et al., 2022)).

For the ACF, the xcorr function in Matlab was used, and then the average distance between peaks of the ACF were used to determine the spatial period. Bootstrapping was applied to determine a statistical cutoff for the peaks, to ensure that there were above noise (previously described in (Biga et al., 2021)).

### 1.8 Statistical tests

All statistical analysis was carried out using GraphPad Prism. For Figure 6C-F, a two-tailed Mann-Whitney U test was used. For Figure 5D&E, a non-parametric One-way ANOVA on ranks (Kruskall-Wallis) followed by Dunn’s multiple comparisons test was used. Significance stars correspond to: ns → p_val_>0.05, * → p_val_≤0.05, ** → p_val_≤0.01, * * * → p_val_≤0.001.

## Supplementary Material

**Figure S1:**
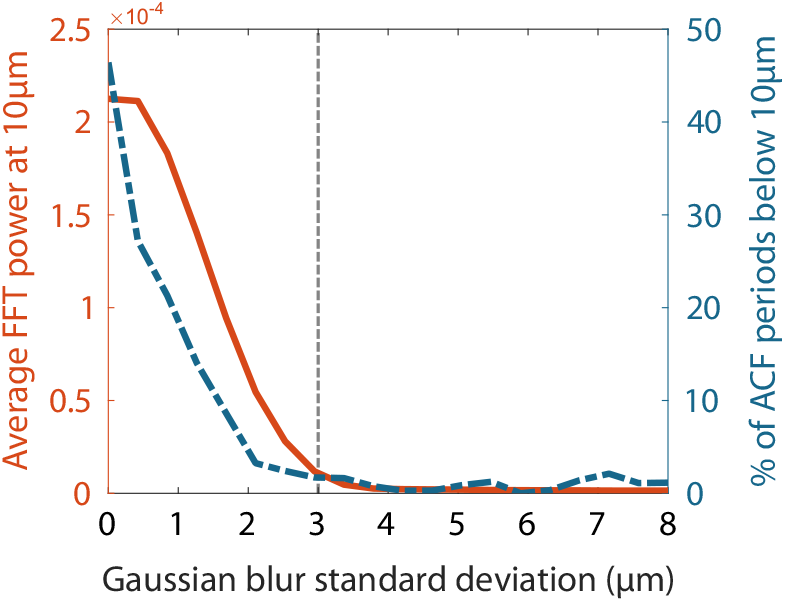
Determining the standard deviation value for the Gaussian blurring applied to the Venus::HES5 fluorescence images in the image analysis pipeline. The aim was to remove contributions of 10μm (the internuclear distance as reported in (Biga et al., 2021)) to the spatial periodicity detected by FFT and ACF. Therefore, we looked at how the Gaussian blur standard deviation affected the power at 10μm in the average FFT power spectrum (all experiments combined). This is shown in the graph in red/solid line. We also examined how the percent of ACF periods below 10μm varied with Gaussian blur standard deviation (blue/dashed line). These both showed a decrease at 3μm (indicated by the dashed grey line), and above this value there was no substantial decrease. Therefore 3μm was chosen as the standard deviation for the blurring step in the imaging pipeline.

**Movie 1:** A 360° view of a whole E10.5 mouse embryo spinal cord slice. Confocal z-stack of fixed Sox2CreERT2^+/−^ mTmG^+/−^. Membrane-tdTomato is in red and tamoxifen-induced membrane-EGFP is in green. The slice shown here is the same as in Figure 2B.

**Movie 2:** A high magnification 360° view of a confocal z-stack of fixed Sox2CreERT2^+/−^ mTmG^+/−^ E10.5 RGCs. The top panel shows the raw membrane-EGFP fluorescence. The middle panel shows membrane-EGFP with marked protrusions (white) overlayed on the central cluster of 3 cells. The bottom panel shows a 3D surface (made using Imaris) of the three central cells, with the three nuclei highlighted in different colours, and with protrusions marked in white.

**Movie 3:** Live imaging of Sox2CreERT2^+/−^ mTmG^+/−^ E10.5 spinal cord slices. The left panel shows membrane-EGFP raw data. The right panel shows the same movie but with tracked protrusions overlayed. Frames were captured every 2 minutes and the time is shown at the bottom right, running to around 4h.

